# Cell painting in activated cells illuminates phenotypic dark space and uncovers novel drug mechanisms of action

**DOI:** 10.1101/2025.05.23.655853

**Authors:** Matylda A Zietek, Akshar Lohith, Derfel Terciano, Beverley M Rabbitts, Aswad Khadilkar, John B MacMillan, R Scott Lokey

**Affiliations:** Department of Chemistry and Biochemistry, University of California, Santa Cruz, 1146 High Street, Santa Cruz, 95064, CA, USA; Department of Biomolecular Engineering, University of California, Santa Cruz, 1146 High Street, Santa Cruz, 95064, CA, USA; Chemical Screening Center, University of California, Santa Cruz, 1146 High Street, Santa Cruz, 95064, CA, USA

**Keywords:** cell painting, phenotypic screening, high-content screening, high-throughput screening, glucocorticoid receptor

## Abstract

As drug and natural product libraries expand, assays for assessing mechanisms of action (MoA) are increasingly critical. Performing cytological profiling using the Cell Painting (CP) assay enables image-based profiling of cellular states upon treatment, yet many bioactive compounds remain uncharacterized due to undetectable cellular effects under standard conditions. To address this, we combined drug dosing with cell activation using the protein kinase C (PKC) agonist phorbol myristate acetate (PMA). Profiling A549 lung cancer cells treated with 8,387 compounds at two concentrations (1 and 10 µM) in both resting and PMA-activated states allowed us to detect phenotypic effects for up to 40% of all screened compounds, effectively illuminating new phenotypic “dark space”. Over 1,000 compounds exhibited phenotypes exclusively under PMA activation, establishing its advantage for MoA studies. We introduce novel quality control measures for CP screens and demonstrate that integrating phenotypic signatures enhances MoA discovery. Notably, 2-methoxycinnamaldehyde clustered with glucocorticoid receptor modulators and induced nuclear translocation, emphasizing the power of this approach in uncovering novel drug mechanisms and, therefore, aiding in improving therapeutic strategies.

## Introduction

A living cell contains myriad complex processes and structures operating concurrently. At any moment, multiple activities coordinate across organelles in an intricately dynamic system. The cell’s interaction landscape depends on conditions such as developmental stage, nutrient availability, and signaling status. Encountering a signaling molecule or stress can induce the cell to undergo physiological and morphological changes—for example, transitioning from a resting to an activated state. This dynamic landscape can be described using various information-rich techniques, including transcriptomics, proteomics, lipidomics, glycomics, and metabolomics. While indispensable for interrogating molecular effects in specific contexts, these techniques do not directly capture a cell’s integrated phenotypic response. An alternative approach is cytological profiling, which uses high-content microscopy to measure hundreds of image-derived features that describe the morphological and functional state of single cells across subcellular compartments, allowing phenotypic effects to be quantified at high throughput ^1,2^. Cell Painting (CP) is a standardized implementation of cytological profiling that facilitates multiplexed fluorescent labeling of key cellular structures, enabling rich phenotypic descriptions across diverse experimental conditions ^3–5^.

CP is used to perform unbiased genetic and chemical screens and has become an assay of choice for exploring mechanism of action (MoA) and target identification studies in drug library screens^6–9^, especially valuable when the target is complex, considered undruggable and/or multipart (polygenic)^10,11^. Uncovering the target(s) or pathway(s) the drug is acting on and, therefore, exposing its mechanism of action is crucial for optimizing drug therapeutic outcomes, increasing efficacy safety, and tackling side effects. CP allows the detection of an immense range of cell phenotypes by its multiparametric approach but becomes an even more powerful technique when coupled with “guilt-by-association” studies. Although discovering the unique cell phenotype upon drug treatment could be extremely exciting, similarities between cell phenotypes treated with different drugs can quickly enhance understanding of the effect of unknown drugs based on similarity to the drug with known MoA. In such an approach, it is standard to use the reference library of drugs with known MoA ^1,2,12^ The bigger and more diverse a reference library is, the better the chance of finding the fingerprint profile of cell features extracted from images similar to the phenotype of interest.

Although Cell Painting is designed as an unbiased, high-content profiling method, it is typically performed on cells maintained in standard culture conditions, where environmental cues, tissue-specific signals, and intercellular interactions are absent. This limitation restricts the assay to the phenotypic landscape of resting cells, in which many signaling pathways remain inactive. As a result, many bioactive compounds fail to produce a detectable phenotype in standard CP screens—particularly when their targets are either not expressed or are functionally silent in the unperturbed state. These limitations give rise to a phenotypic dark space, in which the mechanisms of action for a substantial fraction of compounds remain obscured. We hypothesized that inducing a defined activation state in cells would expand the range of observable phenotypes and help illuminate this dark space. We previously demonstrated the utility of this approach in macrophages (RAW264.7), where lipopolysaccharide (LPS)-induced activation enabled the discovery of novel anti-inflammatory compounds using CP ^13^. Here, we extend this strategy to A549 adenocarcinoma epithelial cells, a widely used model in cancer biology and drug discovery ^9^. A549 cells are also a standard in Cell Painting assays, with extensive reference datasets available for benchmarking^14,15^. In our study, we profiled ∼8,500 known bioactive compounds from TargetMol at two concentrations (1 and 10 µM) to balance sensitivity with cytotoxicity. We performed CP on cells in both resting and activated states, the latter induced by 4-hour treatment with phorbol 12-myristate 13-acetate (PMA), a potent protein kinase C (PKC) agonist that broadly stimulates signaling pathways controlling proliferation, differentiation, apoptosis, and immune response^16,17,18^. Due to its extensive impact on cellular signaling and morphology, PMA is ideally suited as a global activator in high-throughput screens. In this study, we demonstrate how cell activation with PMA reveals previously undetectable phenotypes and enables the functional annotation of compounds that would otherwise remain in the phenotypic dark space. Studying cells treated with our library of bioactives in two concentrations in resting and activated states offers ∼34,000 snapshots of cell “life”, enabling cell network exploration and investigating particular areas of interest or compounds.

## Results

### Drug-exposed A549 cells show the most distinguishable phenotypes in CP when activated with PMA for 4 h and stained according to the JUMP Consortium protocol 3.0

In preparation for high-throughput screening (HTS) of a large compound library, we aimed to establish robust cell-based assay conditions that would allow us to observe distinguishable phenotypes with high confidence. To achieve this, we tested the effects of two broad-spectrum activators on A549 cells: phorbol 12-myristate 13-acetate (PMA) at concentrations of 50 nM and 100 nM and Epidermal Growth Factor (EGF) at 50 ng/mL. These treatments were applied over three different incubation times—28 hours (dosed simultaneously with compounds), 18 h, and 4 h incubation time before cell staining. The CP assay was performed using two sets of dyes, the JUMP Consortium stain set (version 3)^14^ and in-house “classical” stain set, which labels DNA, the Golgi apparatus, F-actin, nuclei, microtubules, mitochondria, and an inflammation marker, nitric oxide synthase (anti-iNOS), as described previously.

Activation of cells with PMA for 4 h resulted in the most distinct phenotype (Fig. 1A), as compared with control (untreated cells), EGF treatment, and other time points. The 100 nM concentration did not provide any additional benefit over 50 nM PMA, therefore, we selected 50 nM PMA for further testing.

**Figure 1.**
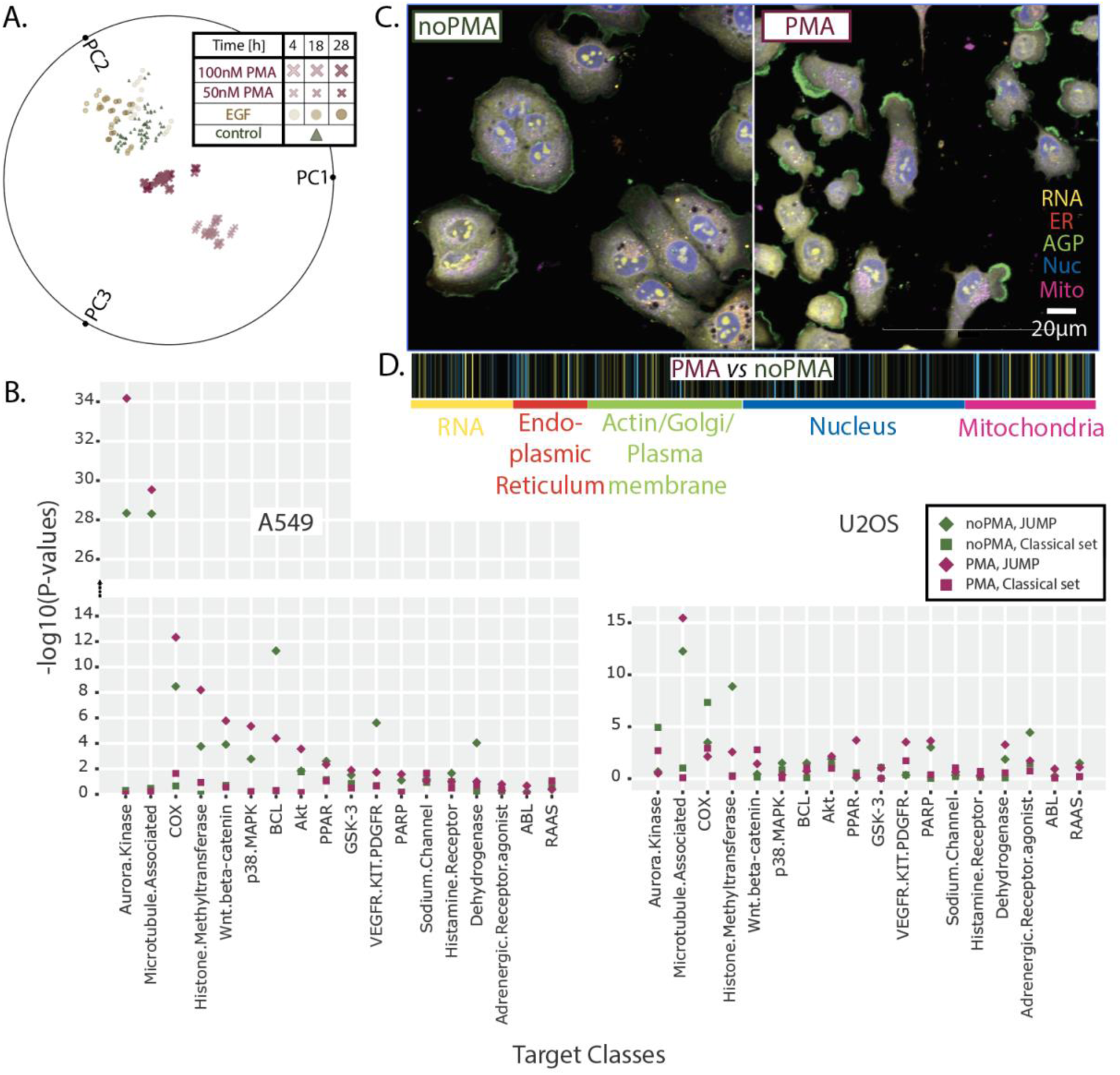
Pilot experiments for High - Throughput, High - Content screen pointed to 4-h activation of A549 cells with 50 nM PMA followed by JUMP staining. **A.** A radial PMA *vs.* EGF PCA plot visualizing how activator, concentration, and time influences the A549 cell phenotype when using our classical stain set. 4-h PMA treatment is causing the most distinguishable phenotype compared to non-treatment. **B.** The scatter plot of p-values from a two-sample, one-sided Kolmogorov–Smirnov (KS) test for a diverse set of target classes shows how the cell’s features could be differently illuminated in different cell lines (A549 *vs.* U2OS) when using different stain sets (JUMP *vs.* classical) and treatments (no PMA *vs.* PMA). A higher -log(p-value) indicates that a particular class is more readily distinguishable from other classes in a specific condition. **C.** Representative images of rested (noPMA) and activated (PMA) A549 cells. The colors in the visualization correspond to different stains in the JUMP stain set used. **D.** The effect of PMA treatment on a cell’s features represented by the fingerprint of 477 cytological features. The mean value for each feature from all PMA control wells was calculated and compared to the mean of respective features from all noPMA control wells; the brighter the color, the more distinguishable the feature is upon PMA treatment.

Subsequently, we performed an experiment to test stain sets for their ability to label informative cell features leading to the best target class separation by using a metric of transformed p-values following a two-sample, one-sided Kolmogorov–Smirnov (KS) test. As the JUMP stain set ^14^ is prevalent in the CP community and is optimized for cost, time efficiency, and ease of use, we tested it alongside our in-house “classical” stain set. This was conducted with two cell lines in parallel. We selected the two lines most used currently in Cell Painting, A549 and U2OS (osteosarcoma cell line with epithelial morphology).

To ensure we can correctly judge the performance of tested conditions, we developed a set of bioactives from 18 different classes (6 compounds representing each class in three replicas, see Supp. Table 1). Half of the set consists of classes that scored well, and half scored moderately in a two-sample, one-sided Kolmogorov–Smirnov (KS) test used in our previous CP study in HeLa cells ^19^. By establishing this compound set from high and medium-scoring classes, we aimed to examine whether newly tested conditions improve scoring for moderately scoring classes but keep well-scoring classes strong. Cells were tested in both resting and PMA-activated cells. When comparing the significance for in-class *vs.* out-of-class target annotation using the KS test (with Pearson correlation as the similarity metric) for tested conditions, we observed the best performance with the JUMP stain set in A549 cells, where 69% of target classes showed significant in-class vs. out-of-class enrichment (Fig.1B). The JUMP stain set showed significant enrichment (*p*<0.05) for 62% of treatments, whereas our classical stain set distinguished only 12% of classes. For the U2OS cell line, 55% of the target classes showed significant enrichment in the JUMP stain set and 22% were enriched significantly in the modified classical stain set.

This study showed that the cell lines are not agnostic with respect to the stain set used. Careful pilot studies should be performed to ensure the best cell line/stain set combination, as some combinations perform very poorly: for example, A549 cells using the classical stain set showed only 2% of classes scoring above the *p*<0.05 threshold (Fig.1B).

Furthermore, we observed that the 4-h, 50 nM PMA treatment affects cells very broadly (Fig.1C), highlighting cellular features from all stained cell structures (Fig.1D), confirming PMA as an appropriate global cell activator triggering multiple cellular pathways.

### A robust Quality Control method increases confidence in HTS CP datasets and decreases screening cost

Generating high-quality datasets from CP HTS requires several QC checkpoints (*ChP*) along the screen pipeline ^4^. Quickly detecting artifacts prevents working with faulty data, lowering cost and labor. An initial checkpoint (*ChP1* Fig.2A) was developed to monitor cell health using live-cell (brightfield) imaging of a few randomly picked plates for each batch, conducted before compound treatment and again before PMA activation. Next, (*ChP2*), before fixation, all 38 control wells per pate (32 negative and 6 positive, Fig.2A, red and blue squares, respectively, in the “Compound dosing to cell culture” panel section) were imaged in brightfield and processed using a machine learning-based linear classification step (Figure 3A). This allowed detection of the level of activation in every plate/well examined and identification of assay plates that had behaved abnormally (showing low activation after PMA treatment or high activation in the carrier, DMSO dosing), thereby allowing such assay plates to be excluded from subsequent steps in the workflow.

**Figure 2.**
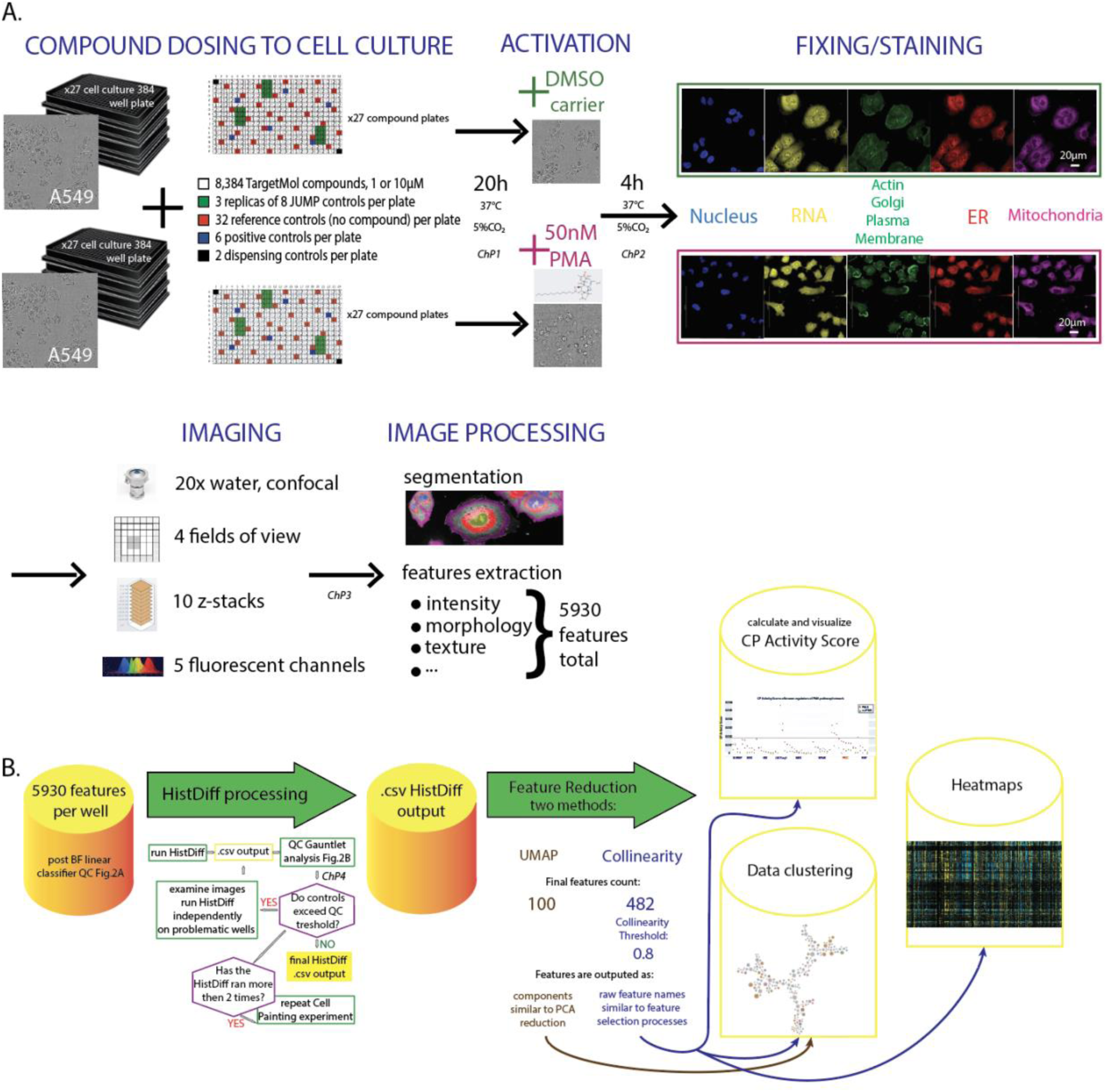
Screen experimental and analytical workflow. **A.** Four sets of 27 384-well plates of A549 cells were processed through a combination of 1, 10 µM TargetMol compound dosing and noPMA vs. PMA treatment. Each plate includes a total of 64 control and 320 experimental wells. Cells were stained using the standard 4-channel JUMP stain set. Imaging was performed using an Opera Phenix high-content screening microscope and image analysis was performed using the microscope’s Harmony and Signal software. *ChP* – CheckPoint for quality control, ER – endoplasmic reticulum **B.** Data analysis pipeline. BF – Brightfield, QC – Quality Control, UMAP - Uniform Manifold Approximation and Projection, PCA – Principal Component Analysis, CP Activity Score – Cytological Profile Activity Score

**Figure 3.**
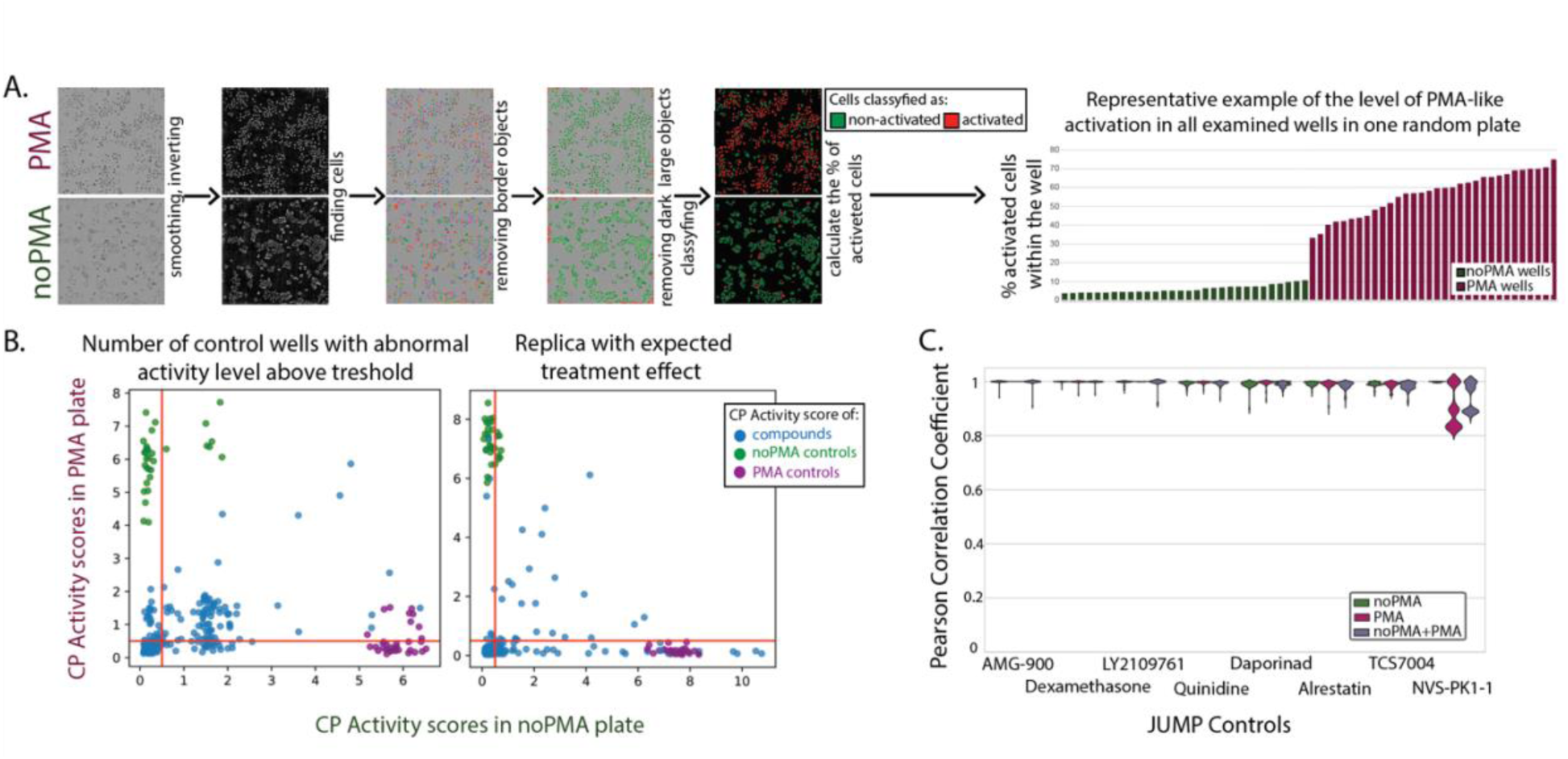
Quality Control checkpoints ensure cost and labor-effective high-quality data acquisition. **A.** To detect if the cell morphology is consistent with the expectation of their appearance as rested or activated, a linear classifier (see Methods) was used on brightfield images. Through the step-by-step image analysis pipeline, the % of activated cells was calculated for each control well. In the shown example on the bottom, wells without activator (noPMA, green) have less than 10% cells behaving like activated ones, whereas most of the wells activated with PMA (maroon) have more than 50% of such cells. **B.** Two examples of plate visualization with QCGauntlet (see Methods). Each dot on the scatter plots represents the CP Activity score for a single well in PMA (Y-axis) *vs.* noPMA (X-axis) plates. The plot on the left was red-flagged as a plate > 22% of all control wells have abnormal activity levels; this plate was assigned to be repeated. On the right is a scatter plot for replica showing the expected overall response to treatment: some compounds show distinguished phenotype just in noPMA condition (higher values on the X-axis), some in just PMA (higher values on the Y-axis), some in both (around diagonal of the plot). There is also dark space represented by compounds in the left-bottom corner of the scatter. **C**. The violin plots of similarity (by Pearson Correlation) of eight JUMP positive controls across the whole screen (within/between plates and across screening batches) in noPMA/PMA/combined conditions.

The next QC checkpoint (*ChP3*) was conducted on the final multichannel, Z-stack CP images. We made an image montage of each 384-well plate and visually examined them to detect any plate artifacts. In case of serious problems, the dataset was discarded and repeated from the cell plating onwards. In a few cases where sections of the plate have obvious technical artifacts like banding or striping, data could be recovered by block normalization, where the large number of reference wells (32) allows for sufficient representation after subdividing the plate, treating the subdivided sections independently.

We then used fingerprints from a single-cell based algorithm (HistDiff) ^2^ to calculate the CP Activity Score, which represents how different each well is compared to the set of reference control wells. This score was used in a QC checkpoint (*ChP4*) for which we developed an application QCGauntlet.py (https://github.com/LokeyLab/QCGauntlet) for visualization facilitating the assessment of overall response to compounds and PMA activator within a plate. QCGauntlet generates various visualizations for CP Activity Scores, with one primary use-case being the generation of scatter plots to compare CP Activity Scores between two different conditions (Fig.3B). We compared the activity of phenotypically active compounds and controls on PMA-treated plates versus untreated plates (Fig.3B). Additionally, the app provides a comprehensive overview of the entire experiment by visualizing CP Activity Scores across all plates. Our study had 27 plates, and the tool allowed us to view the CP Activity Scores of all these plates in a single, consolidated figure consisting of scatter plots for each individual plate, allowing identification of potential batch effects or plate anomalies, thus ensuring data integrity and reliability.

Unfortunately, due to the cost and complexity of CP and related approaches, replicates of the whole screen are not feasible. Instead, one can use the set of the same compounds throughout all plates in a screen, such as the eight maximally diverse positive controls proposed by the JUMP consortium (https://github.com/jump-cellpainting/JUMP-Target?tab=readme-ov-file#positive-control-compounds {JUMP consortium GitHub page}). We used a modified set of these controls (Supp. Table 2) in a 10 µM screen in a randomized position on a plate in triplicates (Fig.2A, green squares in the “Compound dosing to cell culture” panel section).

The similarity of cell response to these compounds was high across replicas within the plate, between plates, and between batches no matter the condition (PMA *vs.* noPMA) (Fig.3C). Such high reproducibility strengthens our confidence in the overall quality of the screen.

### Cell activation expands the phenotypic space illuminated by Cell Painting, providing insights into novel and distinct pathways

To test how much additional value activating cells have on resolving different MoA, we screened a diverse library of bioactives. The 8,387 compounds (TargetMol L4000-CUST, see Supp.SourceData) representing 815 drug classes were dosed in A549 cells for a 24-h incubation (Fig.2A). As this chemical library was tested in two drug concentrations (1 and 10 µM) in resting and PMA-activated cells, and several plates were tested in replicas for quality control (QC) reasons, the combined number of conditions and replicas tested sum up to almost 50,000 individual wells.

The resulting phenotypic fingerprints consist of hundreds of cell features extracted from the segmented regions of the images for each cell and further processed using the HistDiff algorithm, which normalizes each feature vs. the collected DMSO controls from the same plate ^2^ (Fig.2B). The fingerprints were clustered to generate heatmaps of compounds (y-axis) *vs.* cellular features (x-axis) (Fig.4A). As expected, drugs dosed at 10 µM effected more dramatic phenotypic changes in cells compared to the 1-µM treatment, as observed on the global and zoomed-in sections of the heatmap (Fig.4A). However, the higher dose also led to a 2-fold increase in the number of cytotoxic compounds (Fig.4B).

**Figure 4.**
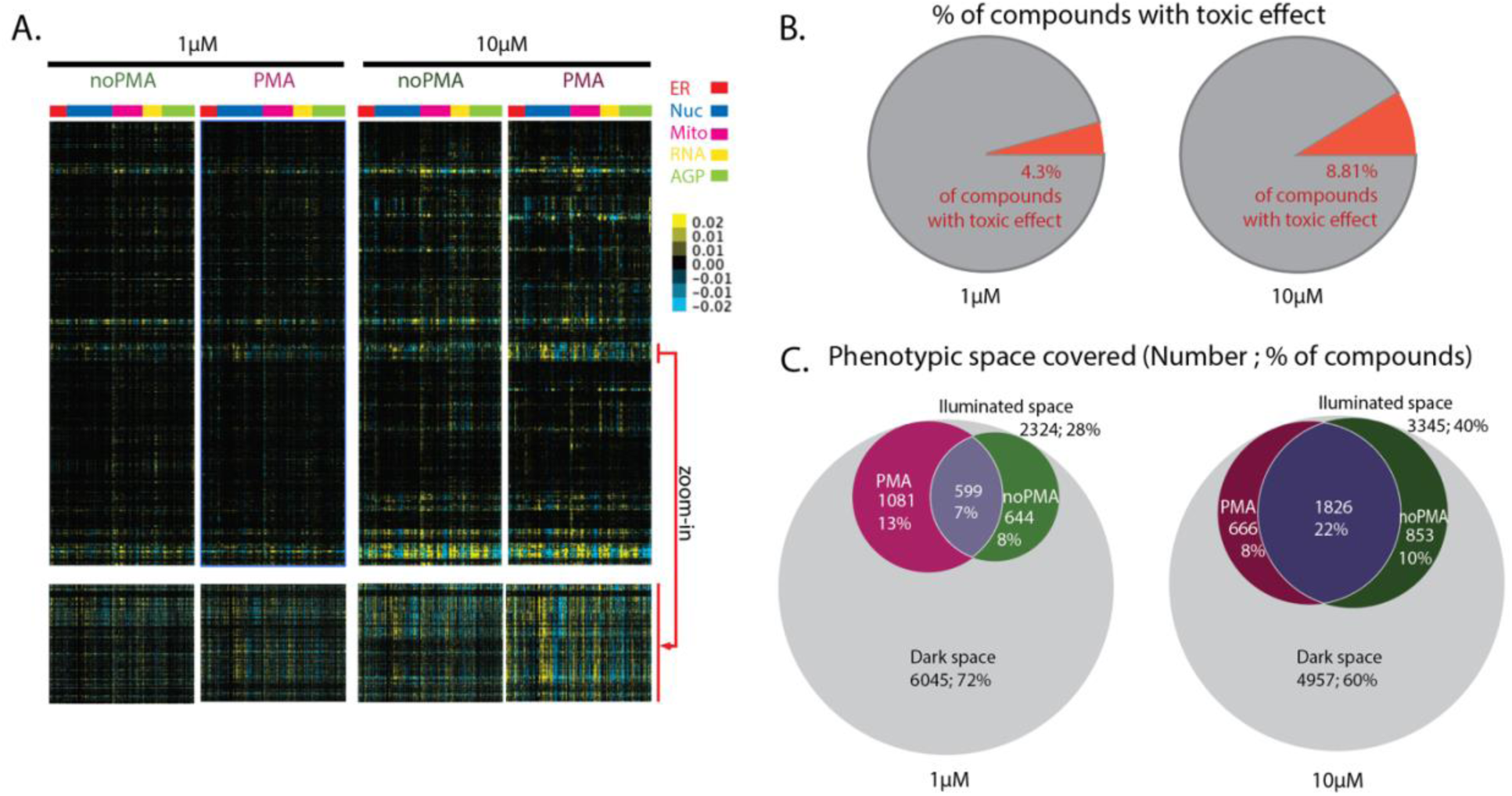
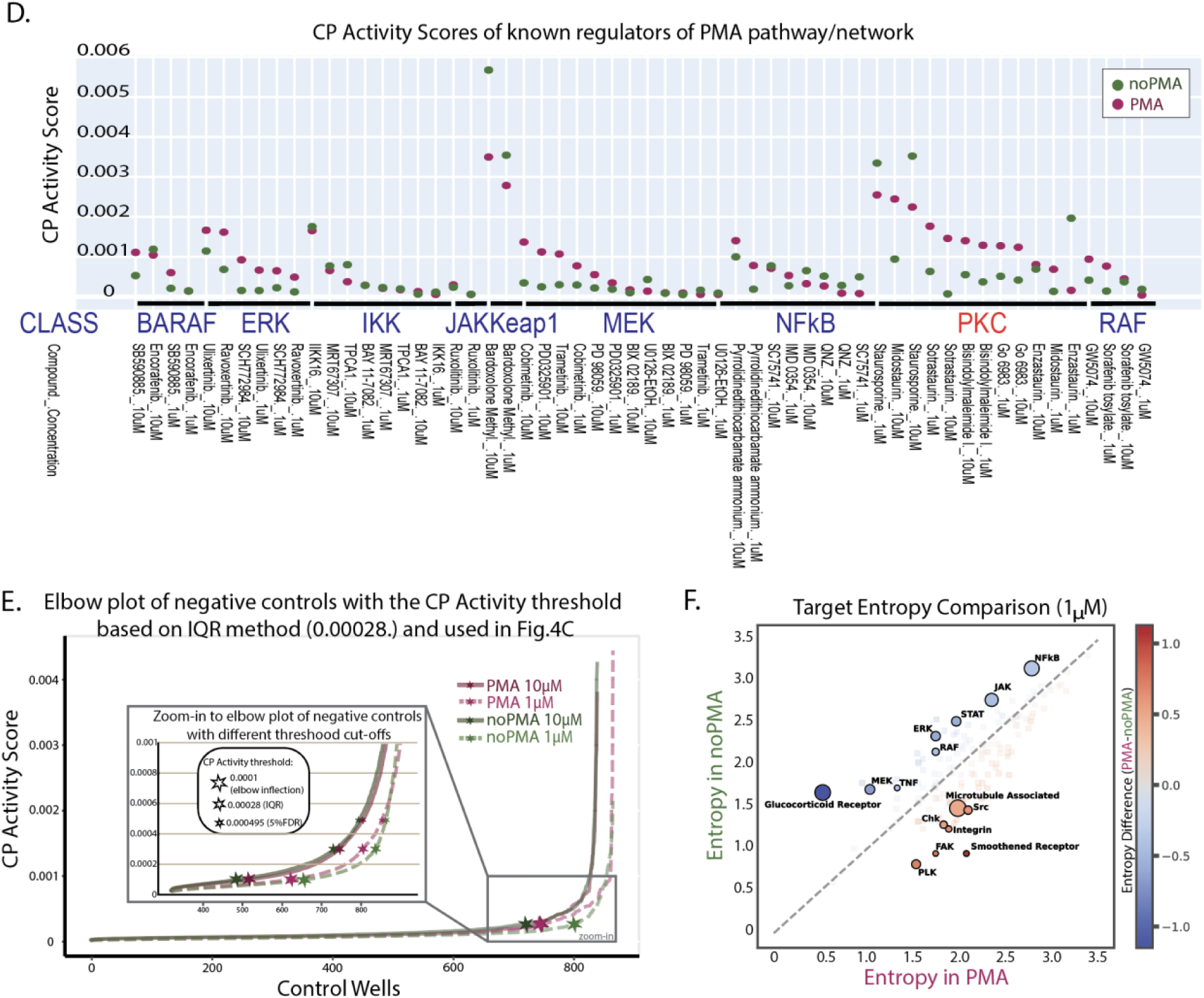
Coupling compound treatment with cell activation reveals new phenotypic space, and this effect is magnified by increasing compound concentration but at the cost of higher cytotoxicity. **A**. The entire heatmap of phenotypic fingerprints representing cell features in noPMA/PMA, 1/10 µM conditions (top heatmap set). The compounds are in the rows and are sorted according to clustering in the 10 µM / PMA condition. The features are in columns and are sorted according to structure stained (ER-endoplasmic reticulum, Nuc-nucleus, Mito-mitochondria, AGP-Actin/Golgi/Plasma membrane) and sorted within a structure group according to clustering in 10 µM PMA condition. The red arrow points to the zoomed-in section of ∼350 compounds (bottom heatmap set). **B.** Treating cells with compounds at 10 µM concentration increases observed cytotoxicity (50% of the average number of cells in control wells) two-fold compared to the 1 µM dosing. **C.** Venn diagrams showing how many / what percentage of compounds are significantly affecting the cells in 1/10 µM screen using the interquartile range (IQR) CP activity score threshold. The maroon color indicates compounds uniquely exerting phenotypic change in activated (PMA) cells, whereas the green color indicates it for resting (noPMA) cells. Compounds that are causing phenotypic changes, no matter the cell’s activation status, are in violet. The effect of a large portion of compounds (grey) is not visible in this particular experimental setup (cell line / condition / compound concentration / stain set used), leaving phenotypic dark space. **D.** The scatter plot of CP Activity Scores of compounds known to affect the PMA network. The effect of the compound on cells is shown in noPMA *vs.* PMA-treated cells. The class to which the compound belongs is indicated in blue, with PKC – the direct target of PMA – indicated in red. **E.** Elbow plot showing the level of phenotypic activity (by CP Activity Score) in control cells (no compound treatment) in different conditions. Cells would display natural heterogeneity in phenotype without any treatment, and there will always be few outliers with significant phenotypic changes (where the elbow raises in a plot). The level of activity in control wells serves to set up what value one judges as cells being active. The stars in the main plot indicate the cut-off established based on the IQR method (see Methods). Inset is a zoom-in to the area of the elbow plot where different cutoff methods for hit calling are indicated by stars. Depending on where on the elbow plot the threshold is set, the ratio between illuminated *vs.* dark phenotypic space will differ (see Supp. Fig.1A, B) **F.** Plot of entropy analysis highlighting targets whose members show greater phenotypic similarity (i.e., become more consolidated, above the diagonal, in blue) upon PMA treatment, vs. targets whose members show decreased phenotypic similarity (i.e. become less consolidated, below the diagonal, in red) upon PMA treatment. Marker size correlates to the number of compounds in that class (Microtubule Associated: 50 members; FAK: 7 members).

The entire dataset (all doses, all conditions, all compounds) is visualized in Fig.4A-C, with the phenotypic features separated by cellular compartment on the heatmaps, enabling global comparison between resting and PMA-activated cells and between the high- and low-dose drug treatments. (Fig.4A, zoom-in).

To take a more detailed approach we calculated the CP Activity Score, which reflects the cumulative effect of treatment on cells and its magnitude (see Methods). Higher CP Activity Score values indicate more features differing more dramatically from no-compound controls, indicating a more robust phenotype.

As a means of quality control, it is of particular interest to focus on PMA’s direct target - protein kinase C (PKC). This enzyme plays a critical role in signal transduction pathways and regulates cell growth and differentiation ^18^. PMA exerts its effect on distinct cell metabolic areas through its intricate network of affected pathways. Indeed, thanks to PMA efficacy, we can clearly observe the phenotypic effect of PKC, MEK, and ERK regulators in PMA-treated cells, among others, giving us confidence in the screen results (Fig.4D).

To assess the impact of our cell activation platform, we sought to evaluate the benefit of illuminating new phenotypes versus the cost of doubling the screen size. To do this, we needed to measure the size of the phenotypic space covered by each condition (resting or PMA-activated, 1 or 10 µM) (Fig.4C). We selected a normalized CP Activity score threshold for distinguishing bioactive from inactive compounds, defining the inactive (“dark”) space using the combined 3404 DMSO control wells (see Fig.2A, red wells). We used the interquartile range (IQR) method to establish the activity threshold (indicated by stars in Fig.4E), in which the outliers above the 75th percentile were excluded since they do not fit the general distribution. Through this approach, we determined a normalized CP Activity Score for the cut-off as 0.00028. Using this threshold, when comparing the different drug concentrations, we could see that 28% and 40% of all screened compounds, in 1 µM and 10 µM dosing, respectively, produced an illuminated effect on cell phenotype (Fig.4C). Since it is expected that a higher dose gives a more pronounced effect (albeit a less informative effect, with more cytotoxic and off-target effects), this result validates our approach. When comparing PMA-activated versus resting cells, as expected, we find that there are some compounds that are illuminated no matter the cell state: 7% of compounds at 1 µM and 22% of compounds at 10 µM dosing (violet shading in Fig.4C).

The most interesting compounds are those that were only illuminated in one of the two conditions - some were specific to resting cells (Fig.4C in green), and some were specific to PMA-activated cells (Fig.4C in maroon). This confirms that cell activation uncovers additional phenotypic space. The 13% of all screened compounds in 1 µM assay would stay in the dark space at the lower dose if the assay did not include PMA activation; this means we now have insight into the MoA for more than 1,000 drugs.

We also present alternate methods for selecting the activity threshold (Supp.Fig.1A, B). We found that the illumination rate varied wildly, from 16.1 to 80.9%, depending on the different thresholds, as well as the concentrations used in a screen. In these cases, the benefit of using PMA can reach as high as 22.5% of added value (Supp.Fig.1B light maroon).

**Supp.Figure 1.**
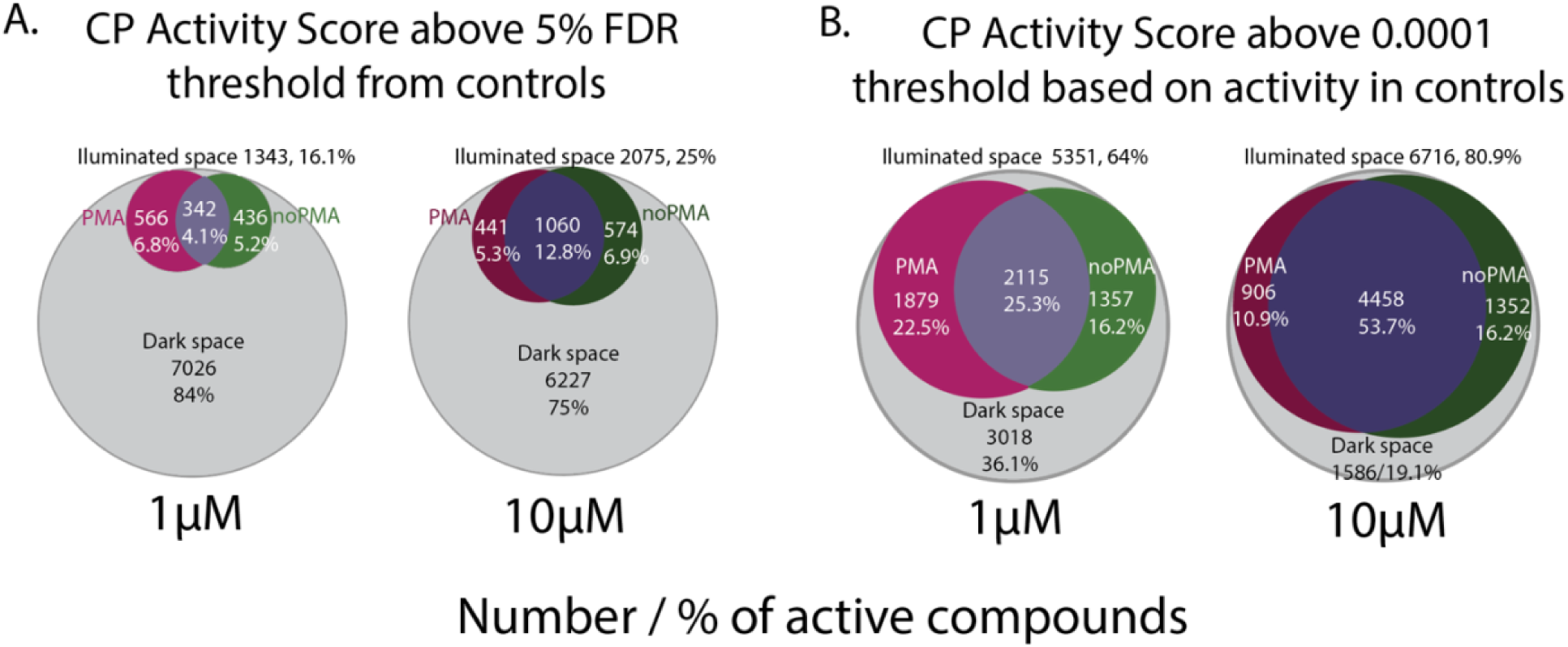
Drawing the cutoff for compound activity is a balancing act between tolerating false positives and accepting false negatives; here, we present a couple of methods in addition to IRQ used in a main text (Fig.4C,E) **A.** The illuminated phenotypic space ranges from 16 to 25%, depending on compound concentration if 5% FDR (False Discovery Rate) hit calling method is used (see Methods and Fig.4E) **B.** If the activity threshold is established just above the inflection point of an elbow plot (0.0001) (see Fig.4E), then the percentage of compounds with significant effect ranges from 64 to 80%

### PMA activation reveals latent pathway structure through target-class consolidation and fragmentation

Extracting the properties of each cell from five channels and multiple cell regions gives rise to thousands (5,930) of features, ranging from signal intensity to morphology and texture. To reduce the dimensions of the dataset, the Uniform Manifold Approximation and Projection (UMAP) technique was employed, narrowing it to the final count of 100 projections (see Fig.2B and Methods); these are output as components similar to PCA reduction ^20^. Based on the UMAP features, a similarity matrix was created and used for clustering with Hierarchical Density-Based Spatial Clustering of Applications with Noise (HDBSCAN, Fig.5A) ^20^. To understand how acute pathway activation alters the functional relationships between compound classes, we examined shifts in phenotypic clustering of inhibitors within their annotated classes in response to PMA activation. Using the pairwise Euclidean distances between the CP fingerprints of the active compounds (using the 0.0001-CP score threshold, Supp.Fig.1B) at 1 µM, we performed entropy analysis to compare their in-class vs. out-of-class distances in the resting vs. PMA-activated condition. This analysis revealed that several target classes, including the glucocorticoid receptor (GR), MAPK pathway components (RAF, MEK, ERK), and JAK/STAT signaling proteins, became markedly more consolidated, that is, they clustered more tightly within their annotated class, upon PMA treatment (Fig.4F, blue). For example, GR-targeting compounds, which displayed more heterogeneous phenotypes in resting cells, clustered tightly in the presence of PMA, consistent with literature describing antagonism between GR signaling and PKC-induced transcription factors such as AP-1 and NF-kappaB ^21,22^. Similarly, inhibitors of RAF, MEK, and ERK coalesced into a shared phenotypic space under PKC activation, likely reflecting their common suppression of the PKC -> RAS -> RAF -> MEK -> ERK cascade, which is robustly activated by PMA ^16^. These observations suggest that PKC activation not only amplifies specific signaling pathways but also synchronizes the phenotypic effects of compounds acting on those pathways, effectively revealing latent pathway-level relationships.

**Figure 5.**
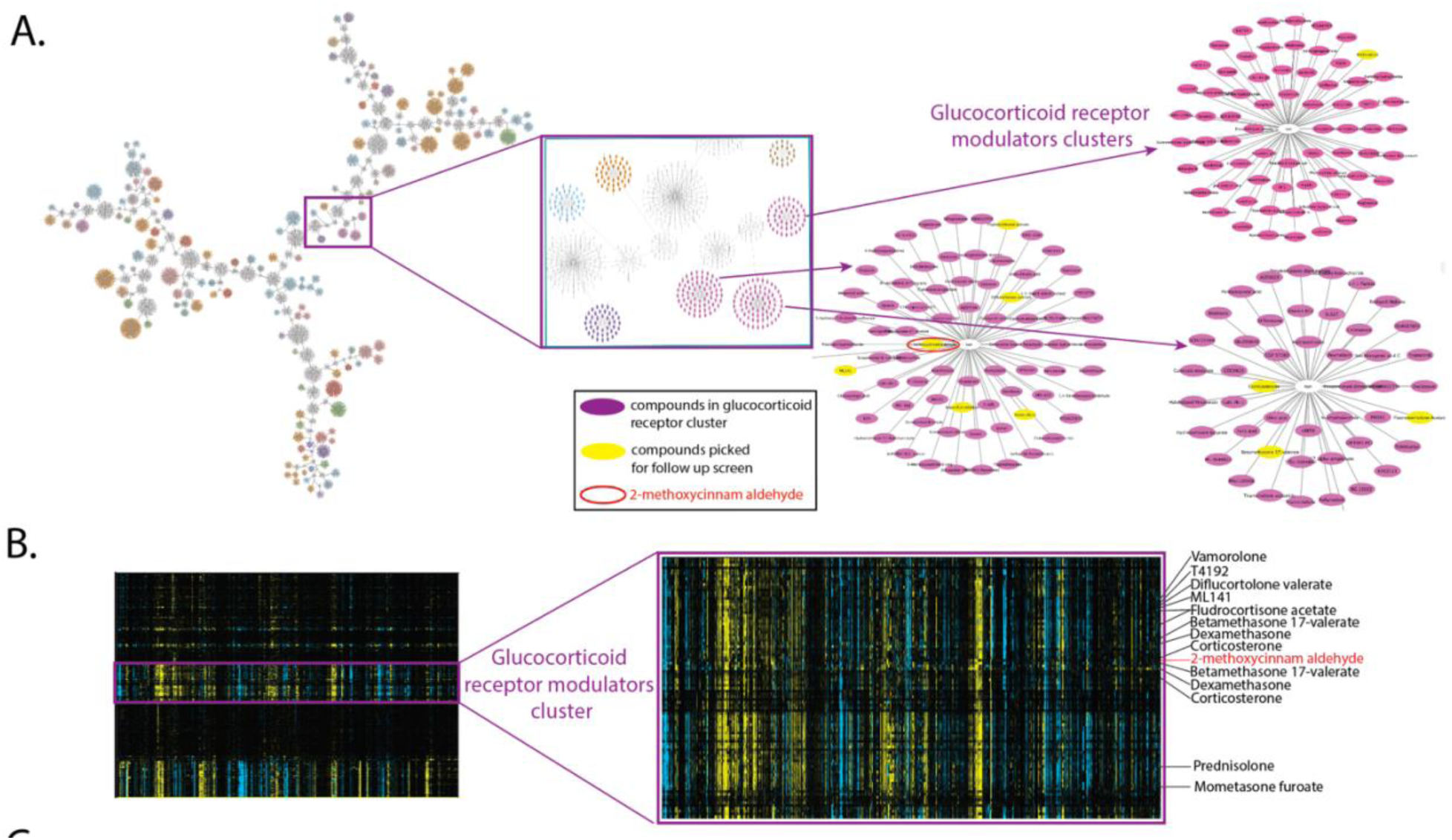
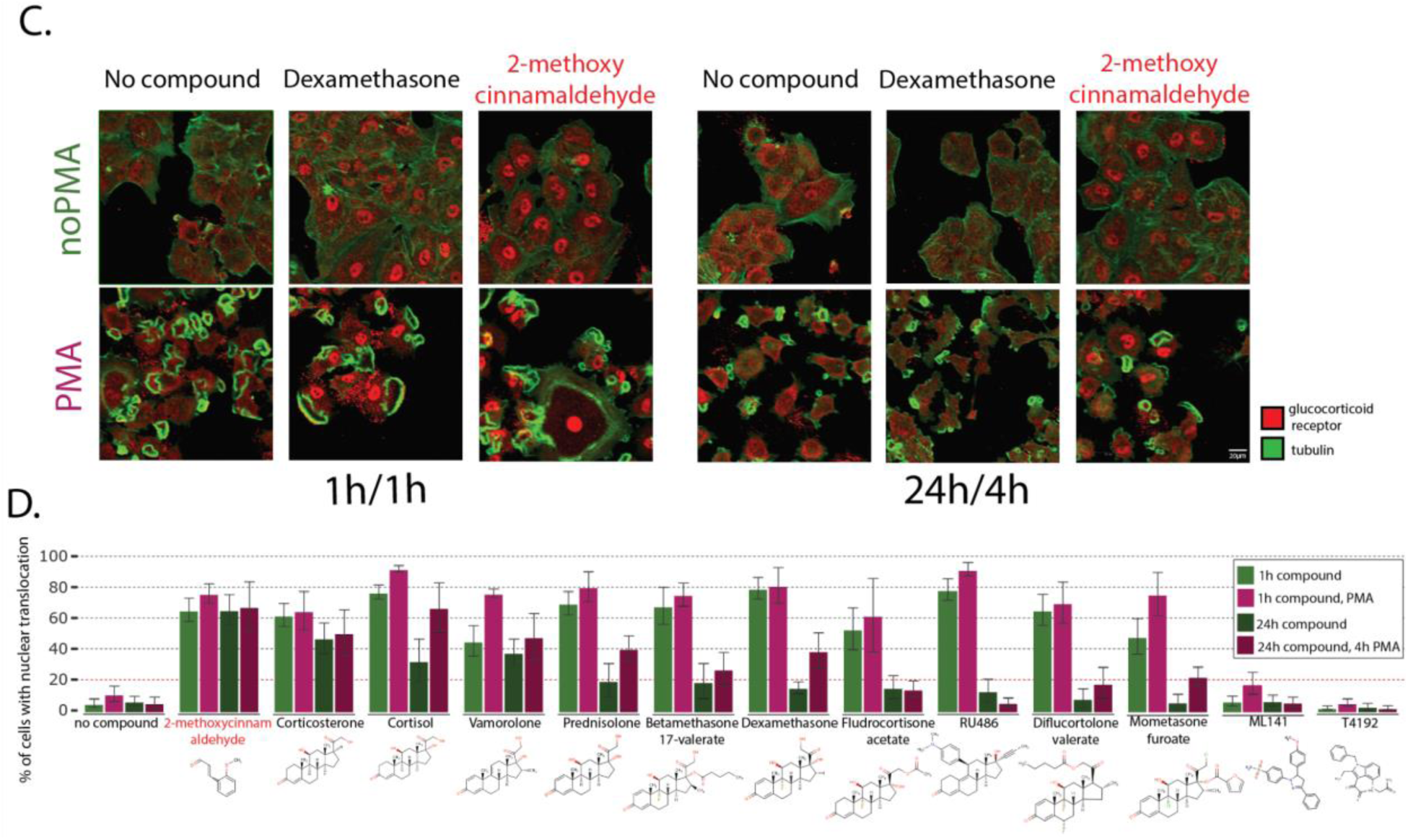
2-methoxycinnamaldehyde clusters with known GR ligands and induces GR nuclear translocation. **A.** The network of clusters of compounds acting on cells in 1/10 µM screen in noPMA/PMA condition based on UMAP features (see Methods). The pink box indicates the area where Glucocorticoid Receptor class compounds are clustering; zooming in allows us to see particular compounds, some of which were picked up for benchmarking (in yellow) **B.** The fragment of a whole screen heatmap of all combined conditions (1/10 µM, noPMA/PMA) with a zoom-in to the cluster of Glucocorticoid Receptor modulators. The cell features are in columns. **C**. The 63x example images of cells treated with no compound / Dexamethasone / 2-methoxycinnamaldehyde for 1 h or 24 h and with noPMA/PMA for 1 h or 4 h in 1 h/1 h and 24 h/4 h treatments, respectively. The tubulin is pseudo-colored green, allowing us to see the cell body, and the glucocorticoid receptor is pseudo-colored red, allowing us to see its varied location in the cytoplasm or nucleus, depending on the treatment. **D**. The nuclear translocation of the Glucocorticoid Receptor was monitored in cells treated with no compound or compounds grouping in the Glucocorticoid receptor cluster. PMA treatment generally enhances nuclear translocation (maroon *vs.* green bars). For some compounds (for example, 2-methoxycinnamaldehyde or cortisol), the nuclear location of receptor persists in 24 h treatment, whereas for others is limited to a shorter, 1 h time frame (for example, RU486 or Diflucortolone valerate). The compound structure is depicted below their names.

In contrast, other target classes became less consolidated under PMA treatment (Fig.4F, red), reflecting greater phenotypic diversity upon PKC activation. Notably, PLK inhibitors, which clustered tightly in the no-PMA condition—likely due to a shared mitotic arrest phenotype— exhibited increased entropy and dispersed into multiple clusters in the activated state. This may result from PKC-induced cell cycle arrest or differentiation, which alters the cellular context in which mitotic kinases are inhibited^23^. A similar loss of consolidation was observed for focal adhesion regulators (e.g., FAK, integrin, Src), whose phenotypes may be modulated by PKC-dependent changes in cytoskeletal organization and adhesion dynamics^24^. Together, these findings suggest that acute pathway activation reorganizes the phenotypic landscape, consolidating the phenotypes of compounds targeting convergent pathways, while dispersing the phenotypes of compounds whose activity becomes context-dependent under activation. This highlights the value of multi-point perturbation—such as combining functional compounds with activators—in revealing hidden structures in phenotypic response space and mapping conditional dependencies between signaling pathways.

### Glucocorticoid receptor agonistic activity predicted from screen network exploration is validated in cells for natural product, 2-methoxycinnamaldehyde

Since compounds targeting GR showed the highest in-class clustering upon PMA treatment among the >800 annotated target classes in the TargetMol library, we examined the compounds in this cluster more closely. Clustering among the known GR modulators were three compounds not previously associated with GR signaling: 2-methoxycinnamaldehyde (T7452, in the “apoptosis” target class), ML141 (T2463, a Cdc42 modulator), and hnps-PLA inhibitor (T4192, a phospholipase inhibitor) (Fig.5A, B). To test these compounds’ ability to modulate GR activity more directly, we employed an immunofluorescence assay that tracks GR nuclear translocation, a hallmark of its activation^25^. The GR normally resides in the cytoplasm, but upon binding to agonists, the complex translocates to the nucleus, where it acts as a transcription factor. As positive controls, we used ten known GR regulators that were also found in this CP cluster. The nuclear translocation is fast and can be observed within 1 h. Because the original CP screen was done using a 24-h compound treatment, we examined the GR nuclear translocation at both 1-h and 24-h drug treatments. We observed that 2-methoxycinnamaldehyde caused GR to translocate from the cytoplasm to the nucleus (Fig.5C red); this effect is clear at 1 h and persists at 24 h. 2-methoxycinnamaldehyde is a relatively simple natural product isolated from the plant genus *Cinnamomum* and in *Illicium verum {*https://lotus.naturalproducts.net/*} {LOTUS database}*), and has anti-inflammatory properties ^26,27^, specifically an inhibitory effect on NF-kappaB ^28^ and antioxidant potential ^29^. Based on the results of the GR translocation assay, one can observe varied persistence of GR nuclear localization. Over 60% of cells treated with 2-methoxycinnamaldehyde retained high nuclear GR levels after 24 h, a proportion greater than for any other compound tested (Fig.5D).

We also investigated the effect of PMA treatment on GR nuclear translocation in the presence of these compounds at both the 1- and 24-h time points (with PMA treatment for 1 h and 4 h, respectively). Interestingly, activating cells with PMA enhanced drug-induced nuclear translocation of GR in almost all cases relative to the no-PMA condition. GR translocation was more pronounced upon PMA treatment at the 24-h time point, consistent with the increased consolidation (decreased entropy) of the GR modulators in the presence of PMA after 24 h. In the presence of PMA, after 24 h 2-methoxycinnamaldehyde was able to induce GR translocation to almost the same level as cortisol.

## Discussion

Cell Painting, or Cytological Profiling (CP), has emerged as a powerful technique for unbiased exploration of cellular phenotypes and drug mechanisms of action. However, the standard approach of profiling resting cells captures only a limited view of the cellular response landscape. In this study, we demonstrate that activating cells with PMA prior to CP analysis substantially expands the observable phenotypic space and enables the detection of otherwise inactive compounds. By screening nearly 8,500 bioactive compounds in both resting and activated states across two concentrations, we were able to illuminate an additional 13% of compounds that showed significant phenotypic effects only in the activated state at 1 µM, representing over 1,000 compounds whose activities would have remained hidden in traditional CP screens.

The choice of activation conditions proved critical for maximizing the information content of our CP profiles. Our systematic comparison of different activators, concentrations, and timing revealed that 4 h treatment with 50 nM PMA provided optimal cellular responses while maintaining cell viability. This finding exemplifies the importance of careful optimization of activation parameters, as too little stimulation fails to reveal new phenotypes, while excessive activation can mask compound effects. The broad impact of PMA on cellular signaling pathways, evidenced by our feature analysis showing effects across multiple cellular compartments, makes it particularly well-suited as an activator for broad screening approaches not directed towards a specific target pathway.

Our implementation of multiple quality control checkpoints, including a novel machine learning-based classifier for detecting cellular activation states quickly from bright-field images, enabled robust and reproducible screening results. The high reproducibility of our JUMP positive controls across plates and batches validates our experimental approach while demonstrating how proper quality control measures can ensure reliable data generation in complex, high-content screens. The development of the QCGauntlet visualization tool further facilitates rapid assessment of plate-level data quality and compound effects, providing a useful resource for the CP community.

The observation that PMA activation drives consolidation of certain target classes while fragmenting others highlights the dynamic interplay between signaling context and compound activity^30^. Pathways such as GR signaling and the MAPK cascade (RAF, MEK, ERK) exhibited tighter clustering upon PKC activation, suggesting that phenotypic responses within these pathways become more uniform when they are strongly engaged or modulated. This is consistent with known antagonistic interactions between PKC-driven transcription factors (such as AP-1 and NF-kappaB) and the GR, where activation of one pathway constrains or homogenizes the functional effects of the other^21^. Similarly, the unification of MAPK inhibitors under PMA likely reflects convergence on the RAS -> RAF -> MEK -> ERK axis downstream of PKC, where inhibition of any node in the pathway blocks a common signaling output ^16^. These results suggest that strong pathway activation can amplify otherwise subtle pharmacologic effects, revealing coherent drug response clusters that might remain hidden under basal conditions.

Conversely, the fragmentation of target classes such as PLK inhibitors, microtubule poisons, and focal adhesion regulators suggests that PKC activation can also introduce heterogeneity, depending on the biological context. PKC-driven changes in cell cycle progression^23^, adhesion dynamics^30^, or cytoskeletal organization likely create new cellular states in which the activities of these inhibitors become more variable. In particular, PLK inhibitors, which induce a stereotyped mitotic arrest in proliferating cells^31,32^, may act less uniformly in cells that have exited the cell cycle or undergone PKC-induced differentiation. Thus, pathway activation not only consolidates phenotypic responses for some target classes but also unmasks biological complexity for others. These findings emphasize the importance of studying compound effects across multiple cellular states to fully capture context-dependent mechanisms of action and uncover latent functional connectivity between pathways.

The biological relevance of activation-dependent phenotypes is demonstrated by our discovery of novel GR translocation upon treatment with 2-methoxycinnamaldehyde. This natural product, previously known for its anti-inflammatory properties, was found to cluster with known GR modulators in our phenotypic analysis, leading to experimental validation of its ability to induce GR nuclear translocation. Notably, this effect was enhanced in PMA-activated cells, suggesting that cellular activation can reveal both new compound activities and potential synergistic effects. The persistent nuclear localization of GR, even after 24 h, upon treatment with 2-methoxycinnamaldehyde, indicates a sustained effect that distinguishes this compound from a panel of classical GR modulators. The presence of an electrophilic aldehyde moiety in this simple compound may indicate that its cellular activity arises from covalent modification of its target, which would be consistent with its prolonged effect on GR nuclear localization.

The decision to screen compounds at both 1 µM and 10 µM concentrations provided complementary insights into compound activities. While the higher concentration revealed more dramatic phenotypic changes and enabled the detection of weakly active compounds, the lower concentration helped identify more specific and potentially physiologically relevant effects while minimizing cytotoxicity. This dual-concentration approach, combined with cell activation, creates a more comprehensive view of compound activities across a broader dynamic range than traditional single-concentration screens in resting cells.

Our findings also emphasize the importance of proper feature selection and data analysis strategies in CP. The comparison of different feature reduction methods - UMAP for similarity-based clustering versus collinearity-based selection for mechanistic interpretation - demonstrates how different analytical approaches can extract complementary insights from the same underlying data. The success of our clustering approach in identifying compounds with similar mechanisms of action, as validated by the GR example, supports the utility of “guilt-by-association” analyses in activated cells for mechanism of action studies.

These results establish activated CP as a powerful approach for expanding the observable phenotypic space in high-content screening. The significant increase in detectable compound activities, combined with the discovery of new mechanisms for known compounds, suggests that many bioactive molecules may have additional activities that are only revealed in activated cellular states. This work provides both a methodological framework and analytical tools for implementing activated CP screens while deepening our understanding of the importance of cellular context in small molecule profiling. Future studies may further expand this approach by exploring different activators and cell types to illuminate additional aspects of compound-cell interactions.

## Materials and Methods

### Chemicals

We obtained a library of 2,035 bioactive compounds (L1700) from SelleckChem, which were used in the pilot. We obtained a library of 8,387 bioactive compounds (L4000-CUST), which were all the known bioactives available in solution format from TargetMol at the time of purchase by CSC (2022-05-03). Daughter plates were prepared with 2.5 mM stocks in 90% DMSO and were used in our CP screen and GR assay. Further details are included in SuppSourceData). These libraries are housed at the UCSC Chemical Screening Center (RRID:SCR\_021114), and more information is available at {https://csc.ucsc.edu/compounds/} {UCSC CSC website}.

### Cell culture

A549 (ATCC Cat. CRM-CCL-185) and U2OS (ATCC Cat. HTB-96) were cultured in DMEM media (Fisher, Cat. 21063029), plus 10% FBS (Fisher, Cat. NC0961964) and 1x penn strep at 37°C, 5% CO2, 95% humidity. For the pilot, CP screen, and GR assay, cells were seeded at 1,250 cells/well in 25 µl by peristaltic bulk dispenser BioTek EL406. For the pilot, we used 384-well Corning 3764 plates, and for the CP screen and GR assay, we used PhenoPlates (Perkin Elmer, Cat. 6057300)

### Drug treatment and activation

All compounds and activators were dosed by Labcyte Echo 650 (Beckman Coulter) and Plate Reformat software, using Labcyte 384PP (Cat. 001-14555) or 384LDV (Cat. 001-12782) source plates.

For the pilot, A549 and U2OS cells were treated with the bioactives from SelleckChem at 10 µM for 28 h. In the CP screen, A549 cells were treated with the 8,387 bioactives from TargetMol at 1 and 10 µM for 24 h. In the 10 µM TargetMol screen, eight positive controls proposed by the JUMP consortium ^9^ with modification (Supp.Table 3) were each used in three replicate wells on each plate. For the GR assay, cells were treated for 1 h or 24 h with 1 µM hit compounds (2-methoxycinnamaldehyde, ML14, or T4192) or known GR agonists (RU486, prednisolone, mometasone furoate, dexamethasone, cortisol, vamorolone, fludrocortisone acetate, diflucortolone valerate, corticosterone, or betamethasone 17-valerate). On each plate, the remaining wells were treated with a DMSO vehicle. There were 64 such wells in the 1 µM screen and 40 wells in the 10 µM screen.

After drug treatment, the cells were activated. For the pilot, cells received 50 or 100 nM PMA (Fisher, Cat. AC356150010) from a stock in 90% DMSO or 50 ng/ml EGF (VWR International LLC, Cat. 10781-692) from a stock in water. They were activated either immediately after drug treatment (which was 28 h before fixing), 18 h, or 4 h before fixing. For the CP screen, 4 h incubation with 50 nM PMA was used, and for the GR assay, 1 h or 4 h incubation with 50 nM PMA was used.

For screening, the 62 control wells (which include reference controls, blocks of JUMP controls, and positive controls) were placed in randomized locations on the assay destination plates, using either the JUMP plate map or a custom randomized plate map developed by Benjamin David and Assistant Professor Paul Jensen, at the University of Michigan Department of Biomedical Engineering using a mixed integer linear program (MILP) performed with a custom design software implemented in Julia version 1.8.2. The exceptions were dispensing controls, wells A01 and P24 reserved for a dye, for visual QC of Echo transfer.

### Quality Control (QC) ChP2: verify activation from brightfield images by a linear classifier

We used an imaging-based, machine-learning linear classifier method to verify that live cells were activated by PMA before proceeding to fix and stain the cells. 2 h after PMA/DMSO treatment (22 h after compound treatment), the entire batch of plates of live cells was moved to random access racks in a LiCONiC StoreX Precision incubator at 37 °C with 5% CO_2. The plates were individually transferred between the incubator and the environmental chamber of the microscope using a PreciseFlex P(3)400 robot with a gripper using PlateWorks scheduling software. A single field of view for each control well was imaged on a Revvity Opera Phenix Plus with a 10x air objective, two peak autofocusing, exposure time of 2ms, transmitted light power at 30%, in widefield mode, and with binning of 2.

An analysis was run online concurrently with imaging in Harmony 5.1 software. The images underwent brightfield correction (a variation of flatfield correction for uneven illumination at the corners), the corrected image was smoothed (using a median filter with a 2 px scale), and the smoothed image was then inverted (using a 60% quantile cut-off). The “find cells” module was run on this inverted image using method B, with a common threshold of 0.21, predicted area >100 \ µm^2, splitting coefficient of 11.6, an individual threshold of 0.16, and contrast ratio >0.04. The cells were then filtered to remove the border objects. For each cell, we calculated 6 standard morphology properties (area, roundness, perimeter, width, length, width to length ratio), brightfield STAR morphology properties (symmetry, threshold compactness, axial, and radial, with profile (4 px), and sliding parabola (curvature of 10), 9 standard brightfield intensity (mean, standard deviation, coefficient of variance, median, sum, maximum, minimum, quantile fraction at 50%, and contrast), and 8 brightfield texture SER properties (scale 0 px and normalized by kernel to locate spot, hole, edge, ridge, valley, saddle, bright, and dark features). Putative cells that were too big (area >2,000 \ µ^2) or too dark (brightfield mean <10,000) were removed from consideration. A linear classifier-based analysis was determined using the training data from a single plate. All the cells in two random resting (no PMA) wells and all the cells in two random PMA-activated wells were the training set of cells for two phenotypes (taking into consideration all the 23 measured features described above for each cell). The goodness of the model was 1.05, with an offset of -1.188. The distinguishing properties selected by machine learning in the final trained model were three texture measurements (dark, valley, and edge) and four intensity measurements (mean, median, contrast, and maximum). This pre-trained classifier was then applied to all cells in all wells of all future plates. In each plate analyzed, the percent activated is calculated (100*a/(a+b) where a is the number of cells that appear activated, and b is the number of cells that appear to have resting phenotype). The percent activated was reported as a bar graph of stimulation (Fig.3A) for each plate. This analysis was performed on all plates from the 10 µM screen. All plates tested passed quality control by the linear classifier, with the criterion that they have no more than two control wells (<10% of the wells) with greater than 15% of the cells therein classified as activated in noPMA wells and have no more than two control wells with less than 30% of the cells classify as activated in the PMA wells.

### Staining

All staining steps were conducted on a BioTek EL406 device equipped with a 10 or 5 μl peristaltic pump cassette and a 192-tube wash module. For the pilot, three stain sets were used on separate plates. Two of the sets, together, are called the “classical set” herein, were stained as described in ^13^. The third set in the pilot and the screen were stained as described by the JUMP consortium in ^14^ (with one assay modification: the drug incubation time was reduced from 48 h to 24 h to limit drug secondary effects). The GR assay was stained with three different stain combinations on one plate. The list of used stains is available in Supp. Table 3. The staining steps are outlined in Supp. Table 4.

### Imaging

For all imaging in this paper, four fields of view for all 384 wells of the plate were imaged on a Revvity (called Perkin Elmer at the time of manufacture) Opera Phenix Plus with a 20x/1.0 NA water objective, two peak autofocusing, in confocal mode with a 50 µm pinhole spinning disk, two simultaneous 2160×2160px cameras, and with binning of 2. A 10-plane Z-stack was collected with 0.8 µm between planes (7.2 µm sample height covered in total). Supp. Table 5 shows a list of channel wavelengths, their laser powers, exposure times, and simultaneous capture pairings used in imaging.

For the screen, the entire batch of fixed, stained cell assay plates was moved to a random-access plate hotel at room temperature. The plates were individually transferred between the rack and the microscope using a PreciseFlex P(3)400 robot with a gripper, using PlateWorks scheduling software. A data transfer to the Signals Image Artist server was run online concurrently with imaging in Harmony 5.1 software.

### Image quantification

For the pilot and the CP screen, image analysis was run using the Signals Image Artist v1.1 web app connected to our Linux data storage server. The images underwent advanced flat-field correction and max projection of the Z-stack. For the 10 µM screen, the images also underwent stitching to create a “global” image, combining all fields of view with dynamic binning. The “find nuclei” module was run on this corrected, max-projected (and stitched in the case of 10 µM) blue channel image using method B, with a common threshold of 0.62, predicted area >30 µ^2, splitting coefficient of 7.0, individual threshold of 0.4, and contrast ratio >0.1. The “find cytoplasm” module was then run on the associated red channel image to find cell regions surrounding the nuclei from the previous step. It used method B, with a common threshold of 0.45 and an individual threshold of 0.15. In the case of the 10 µM screen data, the cells were then filtered to remove the border objects. For each cell in each well, we divided the cytoplasm into three sections: the outermost 30% of the area adjacent to the perimeter is one region, and the rest divided 50% into an innermost area immediately surrounding the nucleus and a third area in between these. We calculated extensive “cell painting properties” of all 5 fluorescence channels with the SER scale set to 2 px. The results table included these cell painting properties for each cell, as well as the object count and all statistics on all properties.

For the GR assay, glucocorticoid receptor translocation was quantified using the following analysis pipeline in Harmony 5.1. It was performed separately for each of the four fields of view in each well. The images underwent a max projection and advanced flatfield correction (to correct for uneven illumination at the corners). The “find nuclei” module was run on the Hoechst image using method B, with a common threshold of 0.62, predicted area >30 µm^2, splitting coefficient of 7, an individual threshold of 0.4, and contrast ratio >0.1. The “find cytoplasm” module was then run using method F, for which we specified the TRITC phalloidin as the “membrane channel” and the AlexaFluor 488 anti-GR as being in the cytoplasm, with an individual threshold of 0.3. The cells were then filtered to remove the border objects. For each cell, we calculated 1 nuclear Hoechst property (intensity sum), 5 nuclear GR properties (intensity mean, median, sum, 50% quantile, and contrast), and 5 cytoplasmic GR properties (intensity mean, median, sum, 50% quantile, and contrast). We then calculated the percent of the cells that could be considered to have nuclear-enriched GR signal, using the following six variables as selection criteria:

- GR nuc sum / Hoechst nuc sum > 0.35
- GR nuc sum / GR Cyto sum > 0.25
- GR nuc mean / GR Cyto mean > 1.7
- GR nuc median / GR Cyto median > 1.7
- GR nuc quantile / GR Cyto quantile > 1.7
- GR nuc contrast / GR Cyto contrast > 0.3

### HistDiff and CP Activity Score

The single-cell quantifications were processed into phenotypic fingerprints composed of a vector of normalized values for each well with a {https://github.com/LokeyLab/HistDiff_standalone} refactored version of the HistDiff algorithm ^2^. Briefly, a histogram of the extracted cell-by-cell values for each feature in each well is constructed and then smoothed and normalized. Following this, the histograms are compared to the corresponding feature histogram constructed with a defined reference well population, evaluating to the dimensionless HisDiff score which describes how different the feature treatment population(s) compares to the reference population.

In a few plates, there were visibly evident stripe/s through the plate where staining was more intense. Since our many reference controls were randomly located throughout the plate, we could divide the plate into datasets, one for the dim rows and another for the bright rows. Each subdivision was processed with HistDiff independently, then concatenated at the end to represent HistDiff fingerprints for the plate.

The CP Activity Score (CP score), which reflects the aggregate strength of the phenotypic fingerprint, was calculated for each condition as follows:

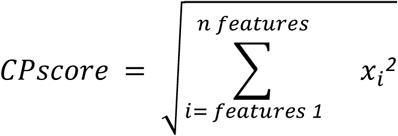

x = feature, n = total number of features

### Feature reduction methods

HistDiff-processed phenotypic fingerprints consisted of 5,880-feature vectors, which we sought to reduce to have a manageable yet informative set of features that still contain most of the original measurement description. Features were selected as described in ^19^, starting with removing uninformative features with zero standard deviation across all treatments. Additionally, using the ‘remove multicollinearity’ function from the Pycaret package (version 3.3), all feature pairs with Pearson correlation coefficients greater than a threshold parameter of 0.8 were flagged, thereby reducing redundancy between the features. Each pair member with the highest mean correlation to all other features was removed, resulting in the 482-feature CP fingerprint for each condition. When looking at the combined view of both conditions (with and without PMA), each set of 482-feature vectors for each compound dose was concatenated on the end, resulting in a 964-feature CP fingerprint for each compound dose in the study. These 964-feature CP fingerprints from collinearity reduction were used for clustered heatmaps and for quantifying the strength of phenotypic response in the study.

We used a different feature reduction method to perform HDBSCAN clustering ^33^ and other pairwise represented analyses. The two (1 µM and 10 µM) sets of the original 5,880-feature HistDiff fingerprints for each treatment condition (compound with or without PMA) were concatenated such that each compound dose treatment had two representatives and processed to 100 dimensions with UMAP ^20^ using the correlation metric and neighbors parameter of 10, with all other parameters left to default. To consider the combined condition for each compound dose, the two (with or without PMA) 5,880-feature fingerprints were concatenated on the end, resulting in an 11,670-feature fingerprint that was processed down to 100 dimensions with UMAP using the same parameters.

### Quality Control ChP4: QCGauntlet.py

We developed a Python tool for CP quality control, which is available in the {https://github.com/LokeyLab/QCGauntlet} {LokeyLab GitHub repository}. This tool, named QCGauntlet, visualizes the CP Activity Scores across the entire screen by generating scatter plots. In these plots, each point represents a single CP Activity Score, with the x-axis corresponding to scores from the PMA condition and the y-axis representing scores from the no PMA condition. Although the tool offers several functions, it was primarily used to display and compare these conditions. To enhance clarity, experimental compounds, and control wells were differentiated using distinct colors and shapes. Two threshold lines—one horizontal and one vertical—were added at the predefined CP Activity Score threshold (see IQR Threshold section), with any score above this threshold classified as active.

The layout of our plate map further informed the expected distribution of controls. For the 1 µM screen, wells with no PMA on the noPMA plate were paired with wells containing PMA on the PMA plate, and *vice versa*. This design resulted in control wells clustering near the origin, while the JUMP control wells appeared in the top right quadrant. In contrast, the 10 µM screen featured reference control locations (no PMA on the noPMA plate paired with PMA on the PMA plate) expected near the origin and positive control wells (PMA on the noPMA plate paired with no PMA on the PMA plate) expected in the top right corner of the scatter plot.

### Quality Control ChP5: after HistDiff and CP Activity Score calculation

In case the control wells were behaving abnormally - having high CP Activity Score (for reference controls) or low CP Activity Score (for positive controls, PMA wells in DMSO plate or DMSO wells in PMA plate), the following scenario was applied: If more than 10% (3 wells) of the control wells score abnormal, then they are removed, and plate normalization scoring is recalculated, excluding those wells from the control population. If there are still 3 or more (10%) control wells with an abnormal activity level, the whole plate is to be rescreened. If a plate has >22% of all the control wells with an abnormal activity level at the outset, the plate is assigned to be repeated.

### Methods for establishing thresholds defining illuminated phenotypic space (pertaining to Fig.4E and Supplementary Fig.1) - IQR, 5% FDR, and elbow inflection methods

Traditionally, CP Activity Score thresholds were determined using the elbow method. In this approach, CP Activity Scores are sorted in ascending order to create an elbow plot, and the inflection point (large stars in the inset of Fig.4E) of this curve is identified as the threshold. The phenotypic space covered using this method is visualized on Supp. Fig.1B.

Alternatively, we explored a False Discovery Rate (FDR) approach. In this method, we evaluated the CP Activity Scores of the reference controls and excluded the top 5 % of controls as outliers. The highest score among the remaining controls was then used as the CP Activity Threshold (small stars in the inset of Fig.4E). This approach is based on the expectation that controls should theoretically exhibit CP Activity Scores near zero since they are not affected by the activator. Therefore, this method emphasizes compounds with the most pronounced stimulation, potentially overlooking those with more subtle activity. Consequently, any compound demonstrating activity beyond that observed in the reference controls is considered active. The phenotypic space covered using this method is visualized on Supp.Fig.1A.

The CP Activity Score threshold for phenotypic activation was established using the interquartile range (IQR) method to exclude outlying control values. Outliers were defined as scores above the third quartile (75th percentile), as these did not conform to the general bell-curve distribution of the control data (see Fig.4E). The highest-scoring control within the acceptable IQR range was then used as the threshold value. This approach ensured that only compounds with CP Activity Scores exceeding this threshold were considered phenotypically active. The phenotypic space covered using this method is visualized in Fig.4C.

### Heatmaps and Clustering

We generated clustered heat maps using a multistep process to further explore the phenotypic signatures derived from our cytological profiling data. First, the fingerprint or feature-reduced raw data were exported as tab-delimited files. We then performed hierarchical clustering on both features and compounds using the BioPython treecluster function. Specifically, we employed the complete linkage method in conjunction with the Pearson correlation coefficient as the distance metric to group similar profiles and reveal underlying patterns.

The resulting clustered data were visualized in Java TreeView, which facilitated interactive exploration and annotation of the heatmaps. This streamlined approach enabled us to efficiently discern the relationships among compounds and cytological features, complementing our statistical assessments with a clear visual representation of the data.

In addition to heatmaps, we conducted a broader clustering analysis by generating a similarity matrix of the fingerprints using the Pearson correlation. The resulting similarity matrix was then subjected to HDBSCAN clustering to identify distinct groups within the data. The clusters obtained from HDBSCAN were subsequently visualized using Cytoscape, with an organic layout applied to the graph to enhance the interpretability of the relationships among compounds and cytological features. This integrated approach allowed us to efficiently discern complex patterns in the data, complementing our statistical assessments with clear visual representations.

## Supp. Method

### Pathway Enrichment Analysis Methodology

#### Data Preprocessing and Filtering

The analysis began with preprocessing of UMAP (Uniform Manifold Approximation and Projection) data from high-content screening of compounds under different treatment conditions (PMA and noPMA) at two concentrations (1 µM and 10 µM). The analysis pipeline consisted of the following steps:

1. Initial Data Selection: The umap_filtering.py script filtered UMAP data from original high-dimensional phenotypic profiles. Compounds were matched with their target annotations from key files containing threshold CP scores above 1e-4, ensuring only compounds with significant activity were included. The data was organized into four primary datasets: PMA 1 µM, PMA 10 µM, noPMA 1 µM, and noPMA 10 µM.
2. Target Filtering: The filter_targets.py script further refined the datasets by keeping only targets represented by at least 3 compounds, excluding the generic “Others” category. This ensured statistical robustness in subsequent analyses by focusing on well-represented targets. The filtering resulted in distinct sets of compounds for each condition, with detailed target distribution statistics saved for reference.

#### Clustering and Entropy Calculation

The entropy analysis was performed using the target_entropy_analysis.py script through the following steps:

1. UMAP-Based Clustering: The script loaded filtered UMAP data for each condition and concentration. For each dataset, two clustering methods were applied:

- K-Means clustering with 50 clusters
- HDBSCAN (Hierarchical Density-Based Spatial Clustering of Applications with Noise) with a minimum cluster size of 10
2. Target Entropy Calculation: For each target across the datasets, Shannon entropy was calculated to quantify the spread of compounds belonging to that target across different clusters. Lower entropy values indicate higher consolidation (compounds are concentrated in fewer clusters), while higher entropy values indicate dispersal (compounds are spread across many clusters). Only targets with at least 5 compounds were included in this analysis to ensure statistical reliability.
3. Entropy Comparison: The script generated entropy comparison plots and data files (entropy_comparison_1uM.csv and entropy_comparison_10uM.csv), which captured the differences in entropy values between PMA and noPMA conditions for each target at each concentration. A negative entropy difference (PMA - noPMA) indicated greater consolidation of a target in the PMA condition, while a positive difference indicated greater consolidation in the noPMA condition.

#### Pathway Enrichment Analysis

The pathway enrichment analysis was conducted using the analyze_pathway_enrichment.py script:

1. Target Categorization: Targets were categorized into specific pathway groups based on biological function using a comprehensive classification system. The classification covered major categories including:

- Neurotransmitter systems (Serotonergic, Dopaminergic, etc.)
- Ion channels and transporters
- Kinase subfamilies
- Nuclear receptor subfamilies
- GPCRs (G-protein coupled receptors)
- Various enzyme types
- Metabolic pathways
- Cytoskeleton and cellular processes
- Cell death and survival mechanisms
- Immune and inflammatory systems
- Epigenetic regulation
- Redox systems
2. Pathway Distribution Analysis: The script analyzed the distribution of pathway categories among targets showing significant entropy differences between conditions. It identified pathways that were preferentially consolidated in either PMA or noPMA conditions.
3. Enrichment Calculation: Enrichment scores were calculated as the ratio of the percentage representation of each pathway category in PMA-consolidated targets versus noPMA-consolidated targets. This quantified which biological systems showed the strongest condition-dependent consolidation effects.

#### Network Visualization

The visualize_cluster_networks.py script was used to create network visualizations that further illustrated the relationships between clusters and targets:

1. Network Construction: Clusters were represented as nodes in a network, with edges connecting clusters based on their proximity in UMAP space. Each cluster was colored based on its most enriched targets, with size proportional to target enrichment.
2. Target Distribution Analysis: For each condition and concentration, target distribution across clusters was analyzed and visualized as heatmaps, providing another perspective on target consolidation patterns.
3. Integrated Visualization: Comprehensive visualizations were created combining UMAP scatter plots, zoomed views of specific clusters, and network representations to highlight the most interesting patterns of target consolidation.

The analysis revealed significant differences in target consolidation patterns between PMA and noPMA conditions, with specific biological pathways showing preferential consolidation in each condition. These findings provided insights into how pathway activation patterns differ under PMA treatment, which activates protein kinase C signaling pathways, and contributes to our understanding of biological system organization in response to different cellular states.

#### Shannon Entropy Calculation

The Shannon entropy for each target’s distribution across clusters was calculated using the following equation:

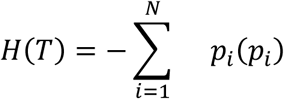

Where:

- *H(T)* is the entropy value for target T
- *p_i_* is the probability of finding a compound with target T in cluster $i$
- *N* is the total number of clusters where the target appears
- *log_2_* indicates the base-2 logarithm, so entropy is measured in bits

In the implementation, the probabilities were calculated by:

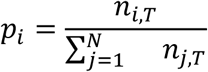

Where:

- *n_i,T_* is the number of compounds with target T in cluster *i*
- The denominator is the total number of compounds with target *T* across all clusters

#### Entropy Difference Calculation

The comparison between PMA and noPMA conditions was quantified as an entropy difference:

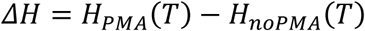

Where:

- *ΔH* is the entropy difference (stored in the ‘Entropy_Diff’ column of the CSV files)
- *H*_*PMA*_(*T*) is the entropy of target *T* in the PMA condition
- *H*_*noPMA*_(*T*) is the entropy of target *T* in the noPMA condition

A negative value of *ΔH* indicates that a target is more consolidated (lower entropy) in the PMA condition compared to noPMA, while a positive value indicates the target is more consolidated in the noPMA condition. The implementation used the entropy function from SciPy’s scipy.stats module, which computes Shannon entropy according to this formula. In the code, this was applied to the normalized distribution of compounds across clusters for each target, with a minimum threshold of 5 compounds per target to ensure statistical reliability.

## Acknowledgment

Thanks to Jon Akutagawa for a fruitful discussion about broad-spectrum cell activators. The instruments used in this research were purchased with the support of the National Institutes of Health grants 1S10RR022455-01A1 and 1S10OD028730-01A1, and these, plus the compound libraries in this study are available through the UCSC Chemical Screening Center RRID:SCR\_021114. We thank Benjamin David and Assistant Professor Paul Jensen at the University of Michigan Department of Biomedical Engineering for providing a randomized plate map. We thank Karolien De Bosscher and Laura Van Moortel at VIB Center for Medical Biotechnology, UGent, for providing the Glucocorticoid Receptor nuclear translocation immunofluorescence assay protocol.

R.S.L. discloses support for the research of this work from the National Institutes of Health (NIH) National Center for Complementary and Integrative Health (NCCIH) and Office of Dietary Supplements (ODS) (NIH-U41AT008718), which is for the Center for High Content Functional Annotation of Natural Products (HiFAN)).

## Notes

### Competing Interest Statement

The authors have declared no competing interest.

