## Supplemental Table 1 for "Cell painting in activated cells illuminates phenotypic dark space and uncovers novel drug mechanisms of action"

Supp. Table 1: SelleckChem compounds used in a cell line/stain set pilot screen.

| Compound | HighScoring Class | Compound | MedScoring Class |
|---|---|---|---|
| **DCC-2036 (Rebastinib)** | ABL | CCT128930 | Akt |
| **GNF-2** | ABL | GDC-0068 | Akt |
| **GNF-5** | ABL | GSK690693 | Akt |
| **GZD824** | ABL | MK-2206 2HCl | Akt |
| **Imatinib Mesylate (STI571)** | ABL | PF-04691502 | Akt |
| **Ponatinib (AP24534)** | ABL | TIC10 | Akt |
| **Dexmedetomidine** | Adrenergic.Receptor.agonist | ABT-263 (Navitoclax) | BCL |
| **Epinephrine Bitartrate** | Adrenergic.Receptor.agonist | ABT-737 | BCL |
| **Isoprenaline HCl** | Adrenergic.Receptor.agonist | AT101 | BCL |
| **Mirabegron** | Adrenergic.Receptor.agonist | Gossypol | BCL |
| **Ritodrine HCl** | Adrenergic.Receptor.agonist | HA14-1 | BCL |
| **Tetrahydrozoline HCl** | Adrenergic.Receptor.agonist | Obatoclax Mesylate (GX15-070) | BCL |
| **CYC116** | Aurora.Kinase | AGI-5198 | Dehydrogenase |
| **Danusertib (PHA-739358)** | Aurora.Kinase | Azathioprine | Dehydrogenase |
| **MK-5108 (VX-689)** | Aurora.Kinase | CPI-613 | Dehydrogenase |
| **MK-8745** | Aurora.Kinase | Gimeracil | Dehydrogenase |
| **MLN8054** | Aurora.Kinase | MK-8245 | Dehydrogenase |
| **PHA-680632** | Aurora.Kinase | Mycophenolate Mofetil | Dehydrogenase |
| **Lornoxicam** | COX | 3-Deazaneplanocin A (DZNeP) | Histone.Methyltransferase |
| **Mesalamine** | COX | Entacapone | Histone.Methyltransferase |
| **Naproxen** | COX | EPZ-6438 | Histone.Methyltransferase |
| **Phenacetin** | COX | EPZ5676 | Histone.Methyltransferase |
| **Sulfasalazine** | COX | MM-102 | Histone.Methyltransferase |
| **Triflusal** | COX | SGC 0946 | Histone.Methyltransferase |
| **1-Azakenpaullone** | GSK-3 | Asiatic Acid | p38.MAPK |
| **AR-A014418** | GSK-3 | LY2228820 | p38.MAPK |
| **AZD1080** | GSK-3 | PH-797804 | p38.MAPK |
| **AZD2858** | GSK-3 | SB202190 (FHPI) | p38.MAPK |
| **BIO** | GSK-3 | SB203580 | p38.MAPK |

| | | | | |
|---|---|---|---|---|
| **CHIR-99021 (CT99021) HCl** | GSK-3 | | Skepinone-L | p38.MAPK |
| **Azacyclonol** | Histamine.Receptor | | AG-14361 | PARP |
| **Azatadine dimaleate** | Histamine.Receptor | | AZD2461 | PARP |
| **Bepotastine Besilate** | Histamine.Receptor | | Iniparib (BSI-201) | PARP |
| **Betahistine 2HCl** | Histamine.Receptor | | INO-1001 | PARP |
| **Brompheniramine hydrogen maleate** | Histamine.Receptor | | ME0328 | PARP |
| **Clemastine Fumarate** | Histamine.Receptor | | Olaparib (AZD2281; Ku-0059436) | PARP |
| **17-AAG (Tanespimycin)** | HSP | | Ciprofibrate | PPAR |
| **AT13387** | HSP | | GW9662 | PPAR |
| **AUY922 (NVP-AUY922)** | HSP | | Rosiglitazone | PPAR |
| **CH5138303** | HSP | | Rosiglitazone HCl | PPAR |
| **Ganetespib (STA-9090)** | HSP | | Rosiglitazone maleate | PPAR |
| **HSP990 (NVP-HSP990)** | HSP | | T0070907 | PPAR |
| **CYT997 (Lexibulin)** | Microtubule.Associated | | Clopamide | Sodium.Channel |
| **Docetaxel** | Microtubule.Associated | | Dibucaine HCl | Sodium.Channel |
| **Epothilone A** | Microtubule.Associated | | Ibutilide Fumarate | Sodium.Channel |
| **Epothilone B (EPO906; Patupilone)** | Microtubule.Associated | | Oxcarbazepine | Sodium.Channel |
| **Nocodazole** | Microtubule.Associated | | Phenytoin sodium | Sodium.Channel |
| **Paclitaxel** | Microtubule.Associated | | Procaine HCl | Sodium.Channel |
| **Cilazapril Monohydrate** | RAAS | | FH535 | Wnt.beta-catenin |
| **Clinofibrate** | RAAS | | IWP-2 | Wnt.beta-catenin |
| **Enalapril Maleate** | RAAS | | IWP-L6 | Wnt.beta-catenin |
| **Moexipril HCl** | RAAS | | | |
| **Olmesartan Medoxomil** | RAAS | | | |
| **Quinapril HCl** | RAAS | | | |
| **Ki8751** | VEGFR.KIT.PDGFR | | | |
| **KRN 633** | VEGFR.KIT.PDGFR | | | |
| **Pazopanib** | VEGFR.KIT.PDGFR | | | |
| **Regorafenib (BAY 73-4506)** | VEGFR.KIT.PDGFR | | | |
| **Telatinib** | VEGFR.KIT.PDGFR | | | |
| **Tivozanib (AV-951)** | VEGFR.KIT.PDGFR | | | |

Supp. Table 2: Positive controls used in 10µM TargetMol screen

| Name | Supplier | Cat # |
|---|---|---|
| **AMG 900** | Cayman Chemical Company | 19176-1 |
| **NVS-PAK1-1** | Cayman Chemical Company | 19964-5 |
| **Alrestatin** | Cayman Chemical Company | 29888-5 |
| **Daporinad (fk866, Apo866)** | Fisher | 501362977 |
| **TCS 7004, Tocris Bioscience** | Fisher | 508810RD |
| **Desametasone, C22H29FO5** | Fisher | MP21993506 |
| **LY2109761** | Sigma | SML2051-5MG |
| **Quinidine** | Spectrum | TCI-Q0006-5G |

Supp. Table 3: List of stains used

| Name | Supplier | Cat # |
|---|---|---|
| **Hoechst 33342** | Life Technologies | H3570 |
| **MitoTracker Deep Red FM** | Invitrogen | M22426 |
| **Glu7-PhalloidinTMR** | (synthetized in house) | |
| **EdU** | | |
| **iNOS (D6B6S) Rabbit mAb** | Cell Signaling Technologies | 13120S |
| **GM130 mouse** | Fisher | BDB610823 |
| **alpha-tubulin-FITC antibody mouse monoclonal** | Sigma | F2168 |
| **Phospho histone H3 (Ser10) Recombinant Rabbit Monoclonal Antibody (9H12L10)** | Invitrogen | 701258 |
| **Chicken anti-Mouse IgG (H+L) Cross-Adsorbed Secondary Antibody Alexa Fluor 488** | Life Technologies | A21200 |
| **Chicken anti-Rabbit IgG (H+L) Cross-Adsorbed Secondary Antibody Alexa Fluor 647** | Life Technologies | A21443 |
| **SYTO14** | Fisher | S7576 |
| **Concanavalin A, Alexa Fluor 488 Conjugate** | Thermo Fisher Scientific | C11252 |
| **Wheat Germ Agglutinin (WGA) Alexa Fluor 555 Conjugate** | Thermo Fisher Scientific | W32464 |
| **Mouse anti-GR G-5, RRID:AB_2687823** | Santa Cruz Biotech | sc-393232 |

Supp. Table 4: Stepwise staining methods

<table>
<tr><th></th><th>CP Screen and Pilot (JUMP stains)</th><th>GR assay</th><th colspan="2">Pilot (“classic” stains)</th></tr>
<tr><td>Live stain(s)</td><td>500 nM MitoTracker Deep Red FM in complete media, 30 min 37°C 5% CO2</td><td colspan="2">Not done</td><td>20 µM EdU and 100 nM MitoTracker Deep Red FM in complete media, 1 h 37°C 5% CO2</td></tr>
<tr><td>Fix</td><td>4% formaldehyde, 6 min RT</td><td colspan="3">4% formaldehyde, 20 min RT</td></tr>
<tr><td>Wash</td><td colspan="4">Twice in PBS</td></tr>
<tr><td>Permeabilize and block</td><td>Not done</td><td>0.1% Triton X-100, 1% BSA, 1 h RT</td><td colspan="2">0.5% Triton X-100, 2% bovine serum albumin in PBS, 0.5 h RT dark</td></tr>
<tr><td>Wash</td><td>Not done</td><td>Twice in cold PBS</td><td colspan="2">Twice in PBS</td></tr>
<tr><td>Click reaction</td><td colspan="3">Not done</td><td>1 mg/mL TMR-azide, 4 mM CuSO4, 2 mg/mL sodium ascorbate in 100 mM Tris</td></tr>
<tr><td>Wash</td><td colspan="3">Not done</td><td>Twice in PBS</td></tr>
</table>

| | | | | |
|---|---|---|---|---|
| **Primary stain** | 8.25 nM TMR-Phalloidin, 5 µg/ml Concanavalin A, 1 µg/ml Hoechst, 1.5 µg/ml WGA, 6 µM Syto14 in 0.1% Triton X-100, 1% BSA 17 min RT dark | mouse anti-GR (and no primary controls) in BSA PBS o/n 4 ℃ | 1:5,000 rabbit anti-histone H3 phospho-S10, 1:1,000 mouse anti-GM-130, 2% BSA in PBS o/n 4 ℃ | 1:5,000 mouse anti-tubulin α, 2% BSA in PBS o/n 4 ℃ |
| **Wash** | Twice in PBS | | | |
| **Secondary stain** | Not done | 2 µM Hoechst, 7 nM TMR-phalloidin, 1:1,000 AF647-anti-rabbit, 1:1,000 AF488-anti-mouse, 2 h RT dark | | 2 µM Hoechst in 2% BSA in PBS 2 h RT dark |
| **Wash** | Not Done | Twice in cold PBS | Twice in PBS | |
| **Mounting medium** | 0.05% Na Az in PBS | | 0.1% Na Az in PBS | |

### Supp. Table 5: Phenix imaging setup

| | Excitation laser and emission filter wavelengths (nm) | Pilot with classic "cyto" stain set 1 | Pilot with classic "EdU" stain set 2 | Pilot with JUMP stain set and CP screen | GR assay |
|---|---|---|---|---|---|
| **B1** | 405, 435-480 | 95% 120 ms | 95% 100 ms | n/a | 95% 100 ms |
| **B2** | 405, 435-550 | n/a | n/a | 95% 160 ms | n/a |
| **G1** | 488, 500-530 | n/a | n/a | 95% 160 ms | n/a |
| **G2** | 488, 500-550 | 95% 500 ms | 95% 200 ms | n/a | 50% 100 ms |
| **Y** | 488, 515-550 | n/a | n/a | 95% 60 ms | n/a |
| **R** | 561, 570-630 | 95% 80 ms | 95% 20 ms | 95% 160 ms | 95% 80 ms |
| **M** | 640, 650-760 | 95% 400 ms | 95% 100 ms | 95% 40 ms (pilot) or 20 ms (screen) | 95% 400 ms |
| **BF** | Brightfield 650-760 nm | 95% 40 ms | 50%, 100 ms | 95% 40 ms | 50% 100 ms |
| **Sequence of capture (camera1/camera2)** | | B1/G2, R/M, BF | | G1, Y, R/M, B1/BF | B1/G2, R/M, BF |
